# RNA sequencing reveals key factors modulating TNFα-stimulated odontoblast-like differentiation of dental pulp stem cells

**DOI:** 10.1101/2025.01.09.632294

**Authors:** Muhammad Irfan, Ji Hyun Kim, Sreelekshmi Sreekumar, Seung Chung

## Abstract

Inflammation is a complex host response to harmful infections or injuries, playing both beneficial and detrimental roles in tissue regeneration. Notably, clinical dentinogenesis associated with caries development occurs within an inflammatory environment. Reparative dentinogenesis is closely linked to intense inflammation, which triggers the recruitment and differentiation of dental pulp stem cells (DPSCs) into the dentin lineage. Understanding how inflammatory responses influence DPSCs is essential for elucidating the mechanisms underlying dentin and pulp regeneration. Given the limited data on this process, a broad approach is employed here to gain a deeper understanding of the complex mechanisms involved and to identify downstream signaling targets. This study aims to investigate the role of inflammation and the complement receptor C5L2 in the odontoblastic differentiation of DPSCs and the associated transcriptomic changes using poly-A RNA sequencing (RNA-seq). RNA-seq techniques provide insight into the transcriptome of a cell, offering higher coverage and greater resolution of its dynamic nature. Following inflammatory stimulation, DPSCs exhibit significantly altered gene profiles, including marked upregulation of key odontogenic genes, highlighting the critical role of inflammation in dentinogenesis. We demonstrate that TNFα-treated odontoblast-like differentiating DPSCs, under C5L2 modulation, exhibit significant differential gene expression and transcriptomic changes. The data presented may provide new avenues for experimental approaches to uncover pathways in dentinogenesis by identifying specific transcription factors and gene profiles.

## 1. Introduction

Dental caries is a common infectious pathology that affects the oral tissues, caused by the invasion of bacteria into the enamel and dentin (1). Current treatment options are limited and often lead to the irreversible loss of pulp tissue (2). The advancement of bacterial caries into dentin prompts dental pulp stem cells (DPSCs) to migrate and differentiate at the site of infection, initiating a regenerative process that leads to the formation of ’tertiary’ dentin (3). This odontoblastic differentiation of DPSCs in response to caries occurs within an inflammatory environment. Inflammation is a crucial immune response that plays a key role in protecting the body from infection and tissue damage. Its complex roles in tissue regeneration, both positive and negative, have gained increasing attention (4). Recent studies have shown that inflammation is instrumental in the regeneration of damaged dental tissues by promoting the recruitment and proliferation of pulp progenitor cells and the differentiation of odontoblasts (5). If this delicate balance between inflammation and regeneration is disrupted, it can lead to irreversible tissue damage. However, there is still limited understanding of the role of inflammation in reparative dentinogenesis and the underlying biology of DPSCs.

Inflammatory cells are recruited to an injury site and release cytokines and growth factors (6). In this study, we utilized the inflammatory cytokine Tumor Necrosis Factor alpha (TNFα), which is known to be elevated in patients with dental caries (7) and plays a central role in coordinating various cellular signaling events (8, 9). Notably, TNFα and ROS, at low levels were reported to promote reparative dentinogenesis (5). TNFα is one of the several pro-inflammatory cytokines involved in bone tissue remodeling and regulation of homeostasis by stimulating osteoclastogenesis and inhibiting osteoblast function. It is also involved in pathogenesis of chronic inflammation and a few inflammatory diseases such as ulcerative colitis and rheumatoid arthritis. However, findings suggests that it also induces osteogenic differentiation (10).

Complement C5a second receptor (C5aR2/C5L2), an alternate C5a receptor, plays a significant role in regeneration and inflammation (11). C5L2-deficient mice exhibited increased mortality, hepatic necrosis, and impaired liver regeneration following partial hepatectomy, emphasizing its critical function in these processes. Our previous study demonstrated that C5L2 modulates brain-derived neurotrophic factor (BDNF) secretion in human dental pulp stem cells (DPSCs). Silencing C5L2 resulted in increased BDNF production, which may accelerate nerve regeneration. This effect is mediated via the p38^MAPKα^ pathway, as shown in our findings (12).

An alternative therapeutic approach involves regenerating dentin and pulp using biomimetic tissue and stem cell engineering. In regenerative medicine, stem cells are being proposed since last decade with attention focused on mesenchymal stem cells (MSCs) as a therapeutic option due to their protective and regenerative abilities (13). Growing evidence are indicating the importance of paracrine signaling induced by MSCs as a supportive mechanism of regeneration of respective damaged tissue (14–16). Currently, there is growing evidence that human DPSCs have many similarities to bone marrow-derived MSCs (BMMSCs) and can be easily isolated, thus having a clear advantage over the costly and invasive techniques required with MSCs collected from bone marrow and DPSCs can be isolated from various dental soft tissues without any invasive technique, such as human dental pulp tissue of permanent teeth (17). In terms of stem cell therapy and regeneration capacity, several advantages and superiority of DPSCs over MSCs have been reported (18, 19). Not only stem cell markers, DPSCs also express dentin and odontoblasts differentiation markers such as dentin sialophosphoprotein (DSPP), dentin matrix protein-1 (DMP-1), alkaline phosphates (ALP) and type I collagen (20). DPSCs can be differentiated by modulation with growth factors, transcriptional factors, extracellular matrix proteins, tissues and cells including: osteoblasts, odontoblasts, neuron cells, and hepatocytes (21). Taken together, these aspects represent DPSCs as promising sources in dentin-pulp regeneration complex.

Earlier, gene expression studies used to rely on low-throughput methods, such as northern blots and quantitative polymerase chain reaction (qPCR), which are limited to measure sole transcripts. Methodologies have been advanced to enable genome-wide quantification of gene expression, or better known as transcriptomics. A high-throughput next-generation sequencing has revolutionized transcriptomics by allowing RNA analysis and understanding of the complex and dynamic nature of the transcriptome (22). In this study, we have investigated the transitional aspects of inflammation, its effects, and underlying mechanisms by examining changes in differential gene profiles and downstream transcriptional factors. Specifically, we analyzed odontoblast-like differentiation of DPSCs treated with TNFα, and with or without C5L2 silencing, through poly-A RNA sequencing to identify key genes and transcription factors involved in the odontogenic differentiation process.

## 2. Materials and methods

### 2.1. Chemicals and reagents

Human DPSCs were purchased from Lonza, Pharma & Biotech (Cat. # PT-5025). MEM-alpha, DMEM, PBS, fetal bovine serum, L-glutamine, and Antibiotic–Antimycotic were procured from Gibco™ Fisher Scientific (Waltham, MA, USA). Poly-D-Lysine coated (BioCoat™, 12 mm) round German glass coverslips were purchased from Corning™ Fisher Scientific (Cat. # 354087; Waltham, MA, USA). Human recombinant TNFα was from Invitrogen, Fisher Scientific (Waltham, MA, USA), and a few other chemicals were from Fisher Chemical (Nazareth, PA, USA). siRNA targeting human C5L2, siRNA control and siRNA Reagent System were purchased from Santa Cruz Biotechnology (Dallas, TX, USA).

### 2.2. Cell culture and treatments

Commercially available human DPSCs, which were guaranteed through 10 population doublings, to express CD105, CD166, CD29, CD90, and CD73, and to not express CD34, CD45, and CD133; were further evaluated by immunocytochemistry in cultures with the STRO-1, a stem cell marker. DPSCs were cultured at 37 °C and 5% CO2 in regular/osteogenic media for 72 h in regular growth media (α MEM containing 10% fetal bovine serum (FBS), 1% L-glutamine), and then cells (a few wells accordingly) were transfected using siRNA Reagent System as previously described (12). Then cells were swapped with dentinogenic media (DMEM containing 20% FBS, 1% L-glutamine and antimycotic/antibiotic i.e., 100 μg/mL streptomycin, 100 U/mL penicillin, supplemented with 100 μg/mL ascorbic acid, 10 mmol/L β-glycerophosphate and 10 mmol/L dexamethasone) for further 7 days differentiation, treated with or without TNFα (10 ng/mL) every three days. All the experiments were conducted with different sets of DPSCs (between 2^nd^ and 4^th^ passages) 3 times, and cell proliferation was measured by counting the total number of cells.

### 2.3. Silencing of C5L2 expression by siRNA

Human DPSCs were grown in 6 well plate culture chamber in 2 mL of free-antibiotic medium up to 70% confluence, then transient transfection with siRNAs was performed using the siRNA Reagent System (sc-45064) according to the manufactureŕs protocol as previously described (12). Cells were incubated at 37°C in a CO_2_ incubator in 1 mL of free-antibiotic and free-serum transfection solution containing a mixture transfection reagent and 40 pmols/mL/well of C5L2 siRNA (sc-105165) or control siRNA, which is a non-targeting siRNA designed as a negative control (sc-37007). After an incubation of 6 hours, 1 mL of medium containing 2 times the normal serum and antibiotics concentration was added in each well without removing the transfection mixture. After 24 hours, the medium was aspirated and replaced with fresh dentinogenic medium and incubated further accordingly as mentioned above.

### 2.4. Total RNA-seq with GO and KEGG enrichment analyses

RNA-seq was performed by LC Sciences from RNA isolated from DPSCs pallet collected after 7-day treatment with or without TNFα, or siC5L2 between passages 2-4. RNA was isolated using the Qiagen RNeasy Mini Kit (79216 Qiagen, Germantown, MD). In short, a Poly(A) RNA sequencing library was prepared following Illumina’s TruSeq-stranded-mRNA sample preparation protocol. RNA integrity was checked with Agilent Technologies 2100 Bioanalyzer. Poly(A) tail-containing mRNAs were purified using oligo-(dT) magnetic beads with two rounds of purification. After purification, poly(A) RNA was fragmented using divalent cation buffer in elevated temperature. The DNA library construction is shown in the following workflow. Quality control analysis and quantification of the sequencing library were performed using Agilent Technologies 2100 Bioanalyzer High Sensitivity DNA Chip. Paired-ended sequencing was performed on Illumina’s NovaSeq 6000 sequencing system. For bioinformatics analysis, in house scripts were used to remove the reads that contained adaptor contamination, low quality bases and undetermined bases (23). Then sequence quality was verified using FastQC (http://www.bioinformatics.babraham.ac.uk/projects/fastqc/). HISAT2 (24) was used to map reads to the genome of ftp://ftp.ensembl.org/pub/release-101/fasta/mus_musculus/dna/. The mapped reads of each sample were assembled using StringTie (25). Then, all transcriptomes were merged to reconstruct a comprehensive transcriptome using perl scripts and gffcompare. After the final transcriptome was generated, StringTie (25) and edgeR (26) was used to estimate the expression levels of all transcripts. StringTie (25) was used to perform expression level for mRNAs by calculating FPKM. The differentially expressed mRNAs were selected with log2 (fold change) > 1 or log2 (fold change) < −1 and with statistical significance (p value < 0.05) by R package edgeR

(26). Four biological replicates were used for the analysis.

### 2.4. Real time PCR

Human DPSCs were cultured in 6-well plate at 5 × 104 cells/well, up to 7 days in dentinogenic media. The total mRNA was extracted using RNeasy Mini Kit (Qiagen, Hilden, Germany) and analyzed using the Fisher Scientific NanoDrop 2000 device. The cDNA samples were analyzed sing the Applied Biosystems SYBR green reagent system, according to the manufacture’s protocol. Primer sequences were purchased from IDT. The expression levels of target genes were standardized by housekeeping gene GAPDH using the 2-ΔΔCt method (27).

### 2.5. Statistical analysis

The statistical analyses were performed on at least 3 independent experiments with duplicates or triplicates, and statistical significance was determined using one-way analysis of variance (ANOVA) followed by post-hoc Dunnett’s test (SAS 9.4) to compare the different treatments and their respective controls (p value of 0.05 or less was considered statistically significant). In addition, the data were also analyzed by Tukey’s test for statistical significance in between the groups.

## 3. Results

### 3.1. Odontoblasts-like differentiating hDPSCs gene profile modulated after TNFα ***treatment***

TNFα is one of the several pro-inflammatory cytokines involved in tissue regeneration and differentiation (10). We evaluated the transcriptional changes in DPSCs odontoblastic differentiation under the effects of TNFα. Differential gene expression characterized by a heat map (Fig. 1A) shows significant differences in upregulation of transcriptional and gene expression verified by volcano graph of differentially expressed genes (DEGs) distribution (Fig. 1B). Biological processes such as immune system regulation, intracellular signal transduction, and cell differentiation were also significantly enriched, indicating a broad impact on cellular functions (Fig. 1C).

**Figure 1.**
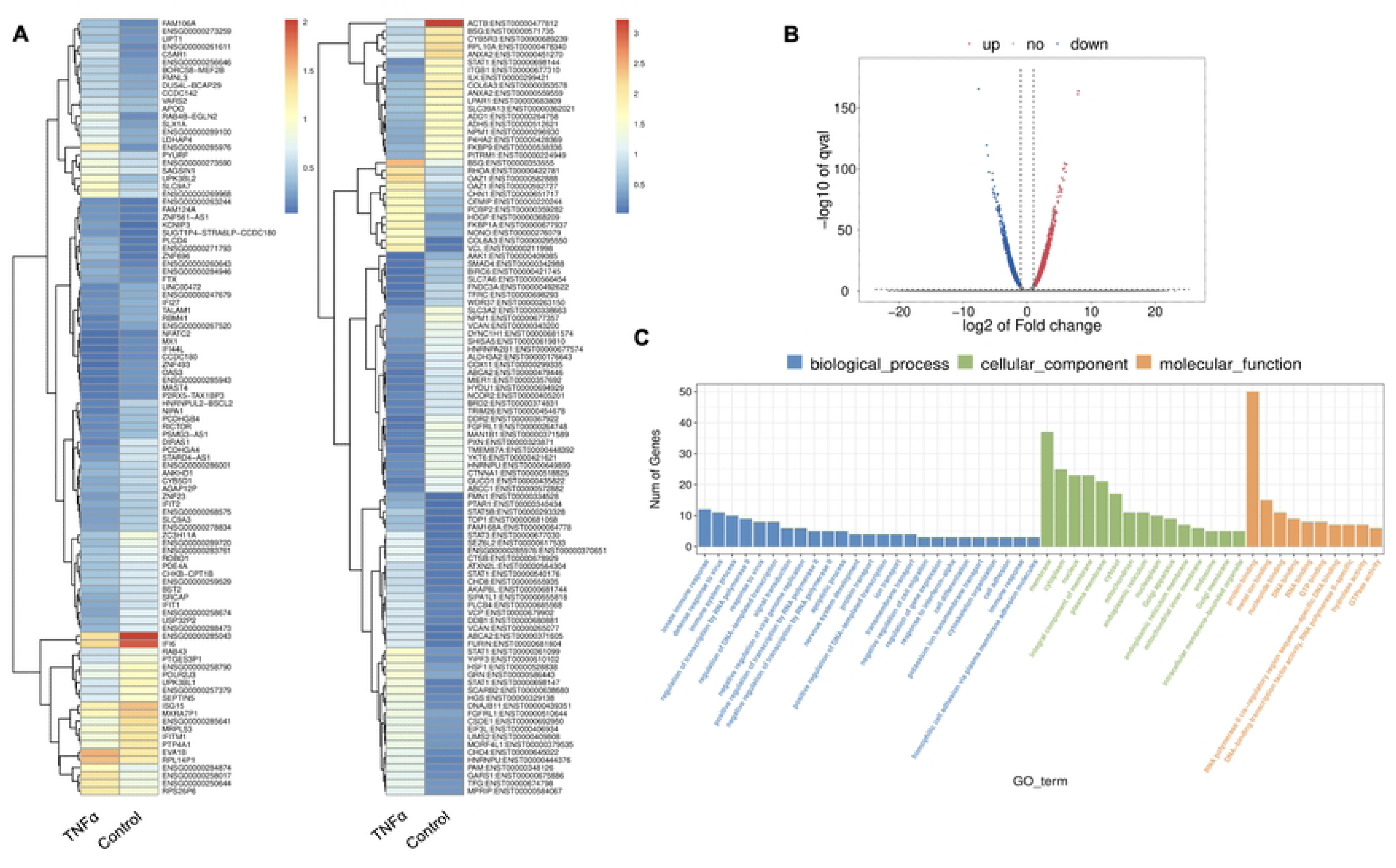
Analysis of differentially expressed genes (DEGs) in RNA-Seq. data. in TNFα treated odontoblast like differentiating DPSCs. **(A**) Heatmap of the differentially expressed genes between untreated or control cells and TNFα hDPSCs in dentinogenic media. Red and yellow stripes in the figure represent high-expression genes, while blue stripes represent low-expression genes. **(B)** Volcano map of differentially expressed genes (DEGs) between control cells and TNFα hDPSCs in dentinogenic media. The x-axis is the log2 scale of the fold change of gene expression in hDPSCs (log2(fold change)). Negative values indicate downregulation; positive values indicate upregulation. The y-axis is the minus log10 scale of the adjusted p values (–log10), which indicate the significant level of expression difference. The blue dots represent significantly upregulated genes with at least twofold change, while the red dots represent significantly downregulated genes with at least twofold change. **(C)** Significant enriched Gene Ontology (GO) terms among control and TNFα treated hDPSCs based on their functions. The top GO terms in the enrichment analysis among biological process, cellular component and molecular function (MF) terms in the enrichment analysis.

Major Gene Ontology (GO) assignments among up-regulated and down-regulated genes include innate immune response and defense response under Biological Process (BP); and integral component of membrane and cytoplasm under Cellular Component (CC); and protein folding and metal ion binding under Molecular Function (MF) [Figure 1. C]. Notably, there was a significant increase in the expression of genes such as Signal transducer and activator of transcription (STAT), Collagen VI (COL6), T-cell factor (TCF), Alkaline Phosphatase (ALP), Dentin Sialophosphoprotein (DSPP), and Dentin matrix acidic phosphoprotein 1 (DMP1). STAT is recognized for its role in regulating inflammation and tissue repair, indicating its potential involvement in reparative dentinogenesis. TCF may influence gene expression associated with odontoblast differentiation and the formation of reparative dentin, where its activation could enhance the regenerative capacity of dental tissues following injury. However, dysregulation of TCF could impair proper dentin formation, leading to developmental defects or compromised repair processes. COL6 is crucial for maintaining the structural integrity of the pulp-dentin complex and supporting odontoblast differentiation and function. Its interactions with other extracellular matrix (ECM) components are likely critical for the mineralization and organization of dentin, contributing to the tooth’s mechanical strength and resilience. ALP is an early marker of odontogenic differentiation in pulp cells, playing a vital role in tissue mineralization and calcification, thus being essential for repair and regeneration processes (28). DSPP and DMP1 work in concert to regulate the formation of reparative dentin; DSPP primarily drives the mineralization process, ensuring the new dentin is sufficiently hard and resilient DSPP (29), while DMP1 plays a key role in organizing the dentin matrix and regulating mineral deposition, ensuring that reparative dentin forms correctly and integrates effectively with existing tooth structure. The upregulation of both DSPP and DMP1 during reparative dentinogenesis underscores their significance in the body’s response to dental injury (30). Collectively, these findings underscore the significant increase in expression of key genes involved in inflammation and regeneration upon TNFα stimulation.

Twenty significant gene ontology (GO) enrichment groups were identified including several enriched pathways related to regulation of immune response (Fig. 2A-B), inflammatory response (Fig.2C) and extracellular structural constituents (Fig.2D). The Sashimi plot illustrates RNA-seq read densities across exons and splice junctions, aligning them with the gene’s isoform structure. Figure 2E presents Sashimi plots for three events on chr14 in TNFα treated cells in dentinogenic media against control (Fig. 2E). For the first gene, there were 21 junction reads in the control samples with inclusion level 1.00. However, in cells exposed to TNFα, the junction reads decreased to 5, with an inclusion level 0.13. This suggests that TNFα exposure induces exon skipping in this particular gene. Similarly, gene 2 and gene 3 exhibited 0.76 and 0.47 inclusion levels respectively following TNFα treatment, whereas control has high inclusion level of 1.00 for both the genes. The difference in inclusion levels between TNFα and control suggests treatment with TNFα results in a decrease in exon inclusion, leading to exon skipping or the production of alternative transcript isoforms. Thus, the results reveal TNFα treatment has a significant impact on the splicing mechanisms of the genes presented, potentially altering their function or expression profiles.

**Figure 2.**
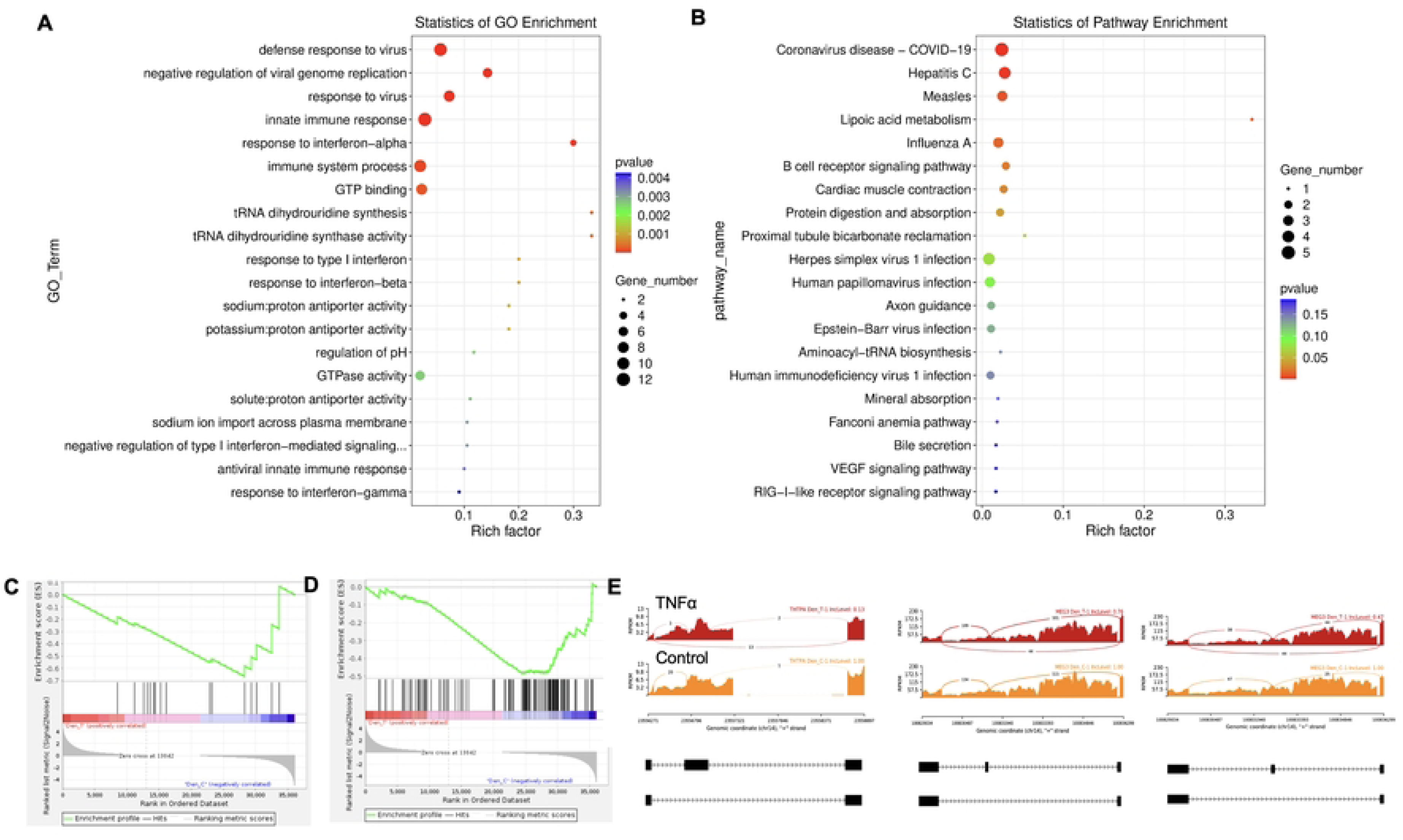
Significant enriched GO terms can be found in TNFα treated hDPSCs in dentinogenic media based on their functions compared with control. (A-B) Go-term and Reactome enrichment pathway analysis of up- and down-regulated DEGs. Dot plot shows top enriched Reactome pathways. The size of the dot is based on gene count enriched in the pathway, and the color of the dot shows the pathway enrichment significance. **(C-D)** Gene set enrichment analysis was performed on DEGs among control and TNFα treated cells in dentinogenic media and found various upregulated and downregulated genes against inflammatory response (C) extracellular matrix structural constituents (D). **(E)** Sashimi plots for quantitative visualization of RNA sequencing read alignments. Data were examined on sashimi plots where it revealed the number of variants and genomic mutation on chr14 in TNFα treated cells in dentinogenic media against control. Red sashimi plots showing variants in TNFα treated group and orange shows in control. While lower black annotations are Read alignments of alternative isoforms and genomic region of interest.

Key transcriptional factors such as lymphoid enhancer factor/ T-cell factor (LEF/TCFs) significantly increased (Fig. 3A) which work through β-catenin pathway to increase BDNF (31), and we have recently reported that enhanced BDNF secretion stimulates DPSCs mediated dentinogenesis (32). TNFα stimulation decreases SMADs activation by inducing NF-κB, which suppresses BMP-2 and TGFβ-mediated SMAD signaling. This suppression disrupts osteoblast differentiation and mineralization, leading to impaired bone formation. The implication is that targeting TNFα or NF-κB could preserve SMAD signaling, potentially enhancing bone mass and offering therapeutic strategies for bone-related disorders like osteoporosis (33). ATF7ip Inhibits Osteoblast Differentiation via Negative Regulation of the Sp7 transcription factor (ref). Also, it is is identified as a crucial regulator of CD8+ T cell immune responses, influencing their effector and memory functions. Mice with a T cell-specific deletion of ATF7ip show increased expression of Il7r and Il2, leading to enhanced CD8+ T cell responses (34).

**Figure 3.**
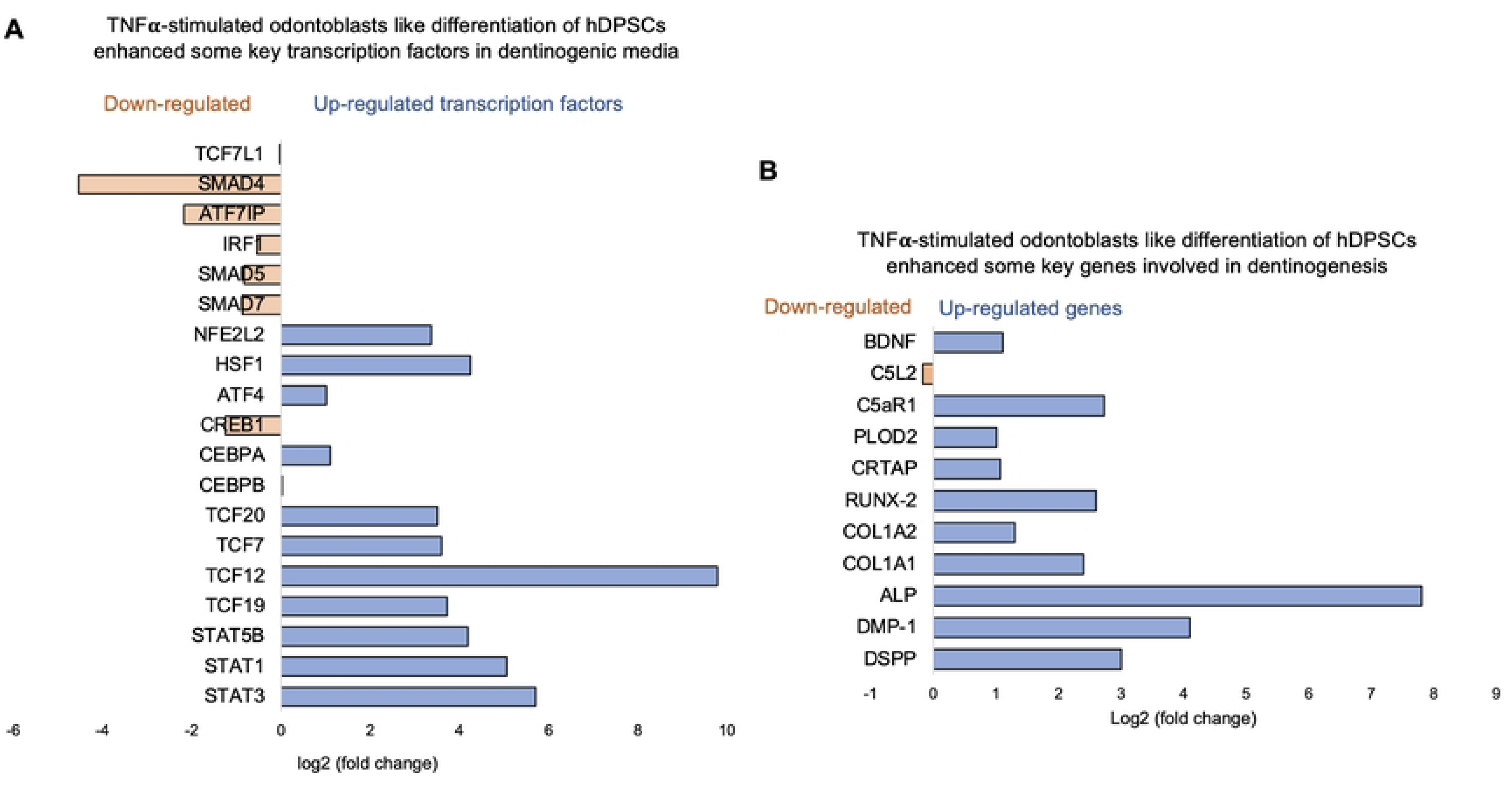
Signaling pathways affected by TNFα in odontoblasts like differentiation of hDPSCs in dentinogenic media. hDPSCs were cultured and treated with or without TNFα for 7 days (twice a week with 3 days interval). Cell lysates were collected, and RNA were prepared using RNeasy mini kit (Qiagen), and next-generation RNA sequencing was done using poly-A-RNA sequencing technique. **(A)** Histogram showing upregulated and activated transcription factors (blue) and repressed or down-regulated transcription factors (orange). It is noteworthy that TCF (7, 12, 19, and 20) especially TCF12 highly up-regulated in TNFα-stimulated odontoblasts like differentiated DPSCs. **(B)** Histogram showing upregulated and activated up-regulated genes (blue) and repressed or down-regulated genes (orange). It is noteworthy that key genes that are involved in dentinogenesis are significantly up-regulated in TNFα-stimulated odontoblasts like differentiated DPSCs.

Similarly, key genes known as odontoblast like differentiation markers, also involved in dentin mineralization such as ALP, DMP-1, RUNX-2 and DSPP (35) were significantly increased (Fig. 3B). The results suggest that TNFα treatment significantly modulates the gene expression profile of odontoblast-like differentiating hDPSCs, particularly impacting immune system regulation, intracellular signaling, and cell differentiation processes. The findings indicate the TNFα’s influence on key transcriptional factors and signaling pathways may disrupt osteoblast differentiation and dentinogenesis, highlighting potential therapeutic targets for enhancing bone formation and treating bone-related disorders.

### 3.2. C5L2 silencing enhances immune regulating and dentinogenic factors

Recent studies, including our own, have identified the activation of the complement system—a key early response to tissue damage—as an additional source of regenerative signals that promote dentin regeneration following carious injuries. We have reported that C5L2 silencing and KO enhances dentinogenesis by regulating BDNF via p38 pathway (12, 36). Here we examined the differential gene expression in siC5L2 treated odontoblastic like differentiating DPSCs. Heat map shows changes in gene expression (Fig. 4A) and volcano graph of DEGs shows upregulated genes against -log10 plot (Fig. 4B); while biological processes such as immune system regulation, signal transduction, and inflammatory response were also significantly enriched, indicating a broad impact on cellular functions (Fig. 4C).

**Figure 4.**
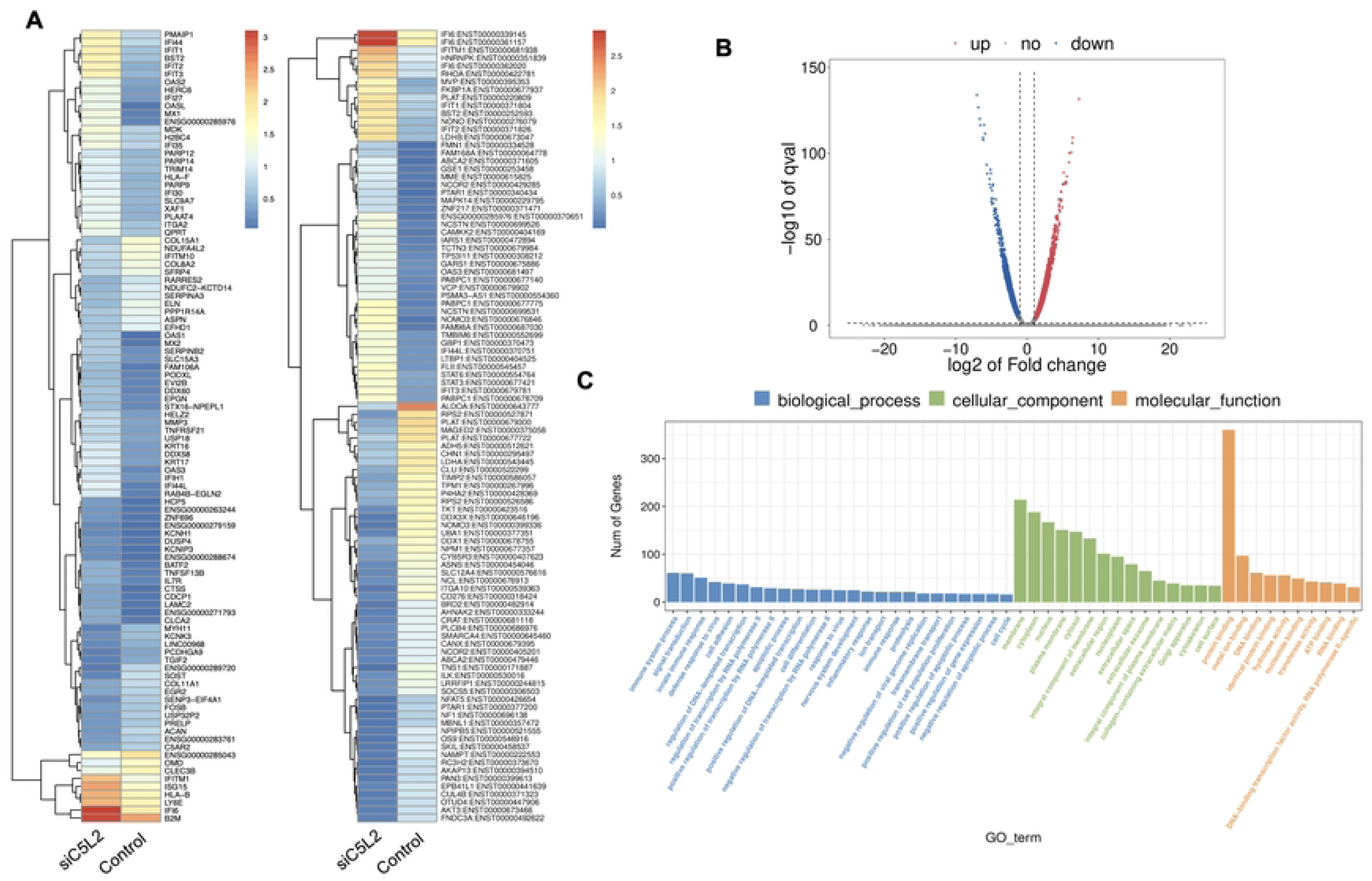
Analysis of differentially expressed genes (DEGs) in RNA-Seq. data. in siC5L2 treated odontoblast like differentiating DPSCs. **(A)** Heatmap of the differentially expressed genes between untreated or control cells and siC5L2 treated hDPSCs in dentinogenic media. Red and yellow stripes in the figure represent high expression genes, while blue stripes represent low expression genes. **(B)** Volcano map of differentially expressed genes (DEGs) between control cells and siC5L2 hDPSCs in dentinogenic media. The x-axis is the log2 scale of the fold change of gene expression in hDPSCs (log2(fold change)). Negative values indicate downregulation; positive values indicate upregulation. The y-axis is the minus log10 scale of the adjusted p values (–log10), which indicate the significant level of expression difference. The blue dots represent significantly upregulated genes with at least twofold change, while the red dots represent significantly downregulated genes with at least twofold change. **(C)** Significant enriched Gene Ontology (GO) terms among control and siC5L2 treated hDPSCs based on their functions.

The major GO terms identified among the genes include innate immune response and signal transduction under Biological Process (BP); integral component of membrane and cytoplasm under Cellular Component (CC); and protein folding and metal ion binding under Molecular Function (MF) [Figure 4. C]. Among these, there was a notable increase in the expression of genes such as Signal Transducer and Activator of Transcription 3 (STAT3), Signal Transducer and Activator of Transcription 6 (STAT6), and Ras homolog family member A (RhoA). STAT3, typically activated in response to inflammation and tissue injury, promotes the survival and proliferation of hDPSCs and odontoblasts. It also facilitates the differentiation of DPSCs into odontoblast-like cells, ensuring that an adequate number of DPSCs are available for the reparative process (37). STAT6 regulates genes involved in extracellular matrix production and remodeling, which are critical for maintaining and repairing dentin. RhoA plays a key role in regulating the cytoskeletal structure of odontoblasts, influencing their migration and positioning during dentinogenesis. Additionally, RhoA signaling is crucial for the differentiation of DPSCs into odontoblast-like cells and impacts the secretion and organization of the dentin matrix. RhoA also helps odontoblasts adjust their cytoskeleton and function in response to mechanical stress, thereby maintaining the integrity of the dentin-pulp complex (38).

Twenty significant GO enrichment groups were identified including several enriched pathways related to regulation of immune response, collagen containing extracellular matrix and cell adhesion (Fig. 5A-B), which are considered critical in regeneration process (39). In caries, innate immune response against bacteria triggered (40, 41) which is important to regulate the oral health. In our results, Enrichment plot and Random ES distribution shows defense response against bacterium have been significantly enhanced after C5L2 silencing (Fig. 5C). Sashimi plots for quantitative visualization of RNA sequencing read alignments. Data were examined on sashimi plots where it revealed the number of variants and genomic mutation on chr16, chr19 and chr11 in siC5L2 treated cells in dentinogenic media against control (Fig. 5D). For LOC, inclusion level (Inclevel) is reported as 1.00, indicating that this splicing or the exon skipping is fully favored in siC5L2. Similarly, gene 2 shows 1 inclusion level following gene silencing, whereas siC5L2 treatment leads to 0.35 inclusion levels in gene 3 against the control. This comparison demonstrates siC5L2 treatment favors exon skipping in two of the three genes shown, leading to a different transcript isoform compared to the control.

**Figure 5.**
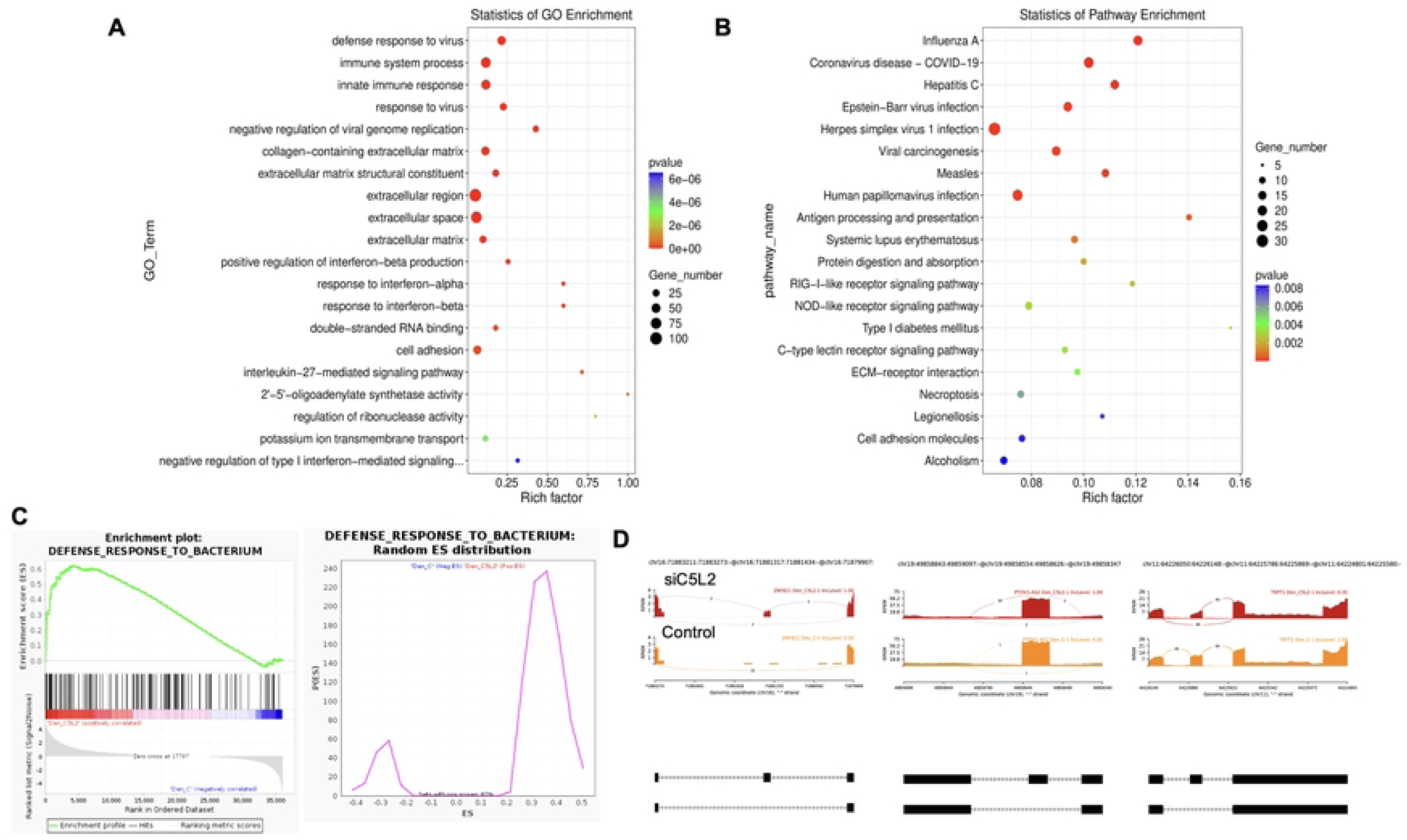
The top GO terms in the enrichment analysis among biological process, cellular component and molecular function (MF) terms in the enrichment analysis in siC5L2 treated odontoblast like differentiating DPSCs. Significant enriched GO terms can be found in TNFα treated hDPSCs in dentinogenic media based on their functions compared with control. **(A-B)** Go-term and Reactome enrichment pathway analysis of up- and down-regulated DEGs. Dot plot shows top enriched Reactome pathways. The size of the dot is based on gene count enriched in the pathway, and the color of the dot shows the pathway enrichment significance. **(C)** Gene set enrichment analysis was performed on DEGs among control and siC5L2 treated cells in dentinogenic media and found various upregulated and downregulated genes against bacterial response. Enrichment plot and Random ES distribution shows defense response against bacterium have been enhanced after C5L2 silencing. **(D)** Sashimi plots for quantitative visualization of RNA sequencing read alignments. Data were examined on sashimi plots where it revealed the number of variants and genomic mutation on chr16, chr19 and chr11 in siC5L2 treated cells in dentinogenic media against control. Red sashimi plots showing variants in siC5L2 treated group and orange shows in control. While lower black annotations are Read alignments of alternative isoforms and genomic region of interest.

### 3.3. Transcription factor 7-like 2 increased by C5L2 silencing

β-catenin and LEF/TCF form a complex with SMAD4 (42), and Silverio et al., (43) proposed that β-catenin and SMAD proteins are needed to activate cementoblast/osteoblast gene expression and promote full differentiation. Bem et al., (44) proposed that alterations in the activity of LEF/TCFs, and particularly of transcription factor 7-like 2 (TCF7L2), result in defects previously associated with neuropsychiatric disorders, including imbalances in neurogenesis and oligodendrogenesis. The canonical Wnt signaling pathway plays an important role in tooth development by pulp capping, enhancing cell proliferation and inducing odontoblast differentiation (45, 46) and accumulated β-catenin translocates into the nucleus and binds to the transcription factor TCF/LEF to promote transcription (47). Again, our results showed that siC5L2 treatment enhanced BDNF secretion related transcriptional factor ultimately contributing to dentinogenesis including TCF/LEF (especially TCF7L2) and SMADs (Fig. 6).

**Figure 6.**
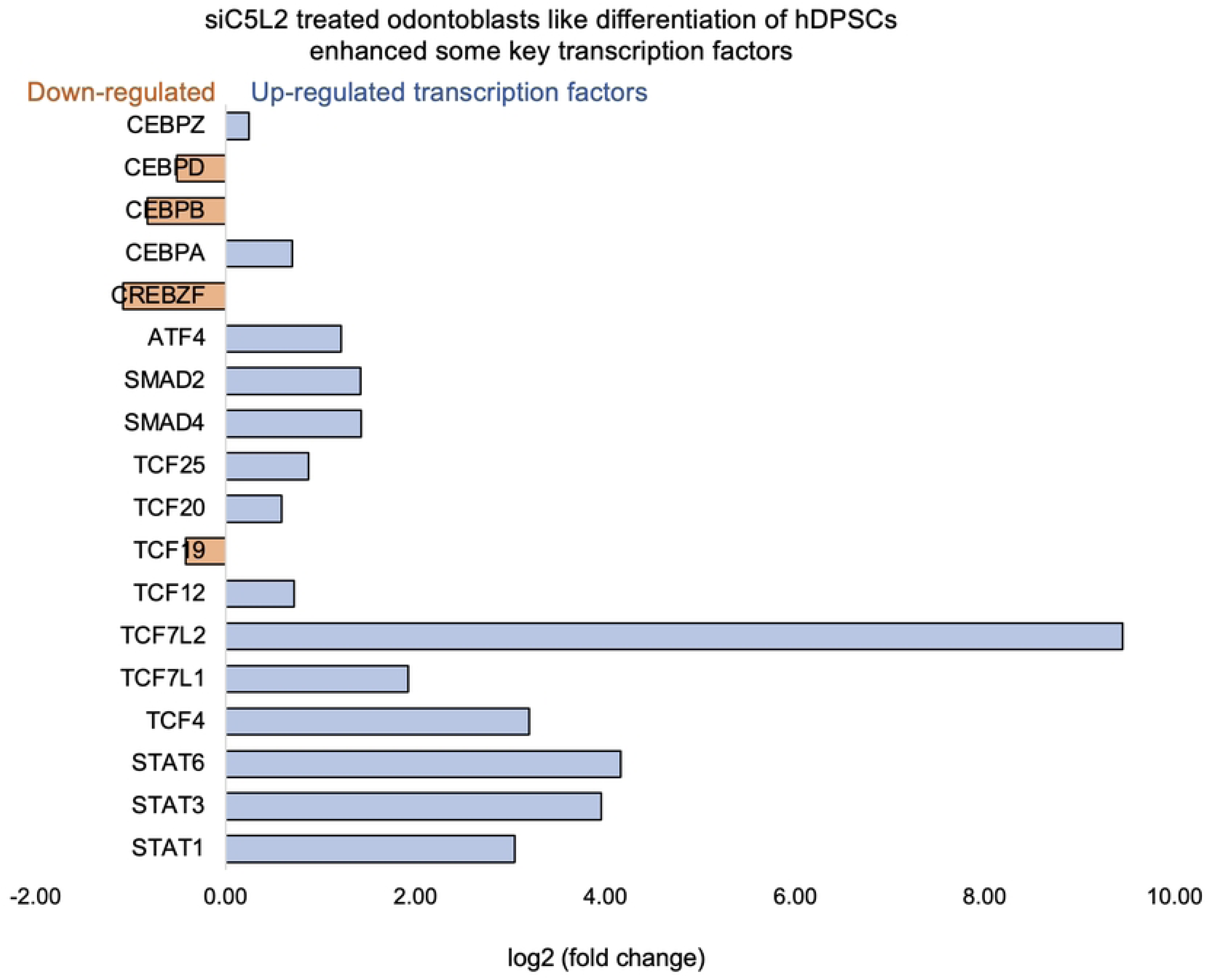
Signaling pathways affected by siC5L2 in odontoblasts like differentiation of hDPSCs in dentinogenic media. hDPSCs were cultured and treated with or without siC5L2 and differentiated for 7 days. Cell lysates were collected, and RNA were prepared using RNeasy mini kit (Qiagen), and next-generation RNA sequencing was done using poly-A-RNA sequencing technique. Histogram showing upregulated and activated transcription factors (blue) and repressed or down-regulated transcription factors (orange). It is noteworthy that TCF (4, 7, 12, 20, and 25) especially TCF7L2 highly up-regulated in siC5L2-stimulated odontoblasts like differentiated DPSCs.

The signal transducer and activator of transcription (STAT) protein family are intracellular transcription factors that mediate many aspects of cellular immunity, proliferation, apoptosis and differentiation. These transcription factors activated through the JAK-STAT pathway, play a crucial role in stem cell proliferation by promoting the transcription of genes associated with cell growth (48). The results suggest that silencing C5L2 increases the expression of transcription factor TCF7L2, which plays a critical role in odontoblast differentiation and dentinogenesis through the Wnt/β-catenin and SMAD signaling pathways. This indicates that targeting C5L2 could enhance these pathways, potentially improving tooth regeneration and repair processes.

### 3.4. Combined effects of TNFα and siC5L2 on immune regulators and dentinogenic markers

Recently, we have reported the synergistic effects of TNFα and siC5L2 treatment on BDNF secretion and dentinogenesis (12, 32, 36). Here, we have examined their effects on differential gene profiling of TNFα and siC5L2 treated odontoblastic like differentiating DPSCs. Heat map shows differential gene expression on both groups (Fig. 7A) while volcano graph of DEGs shows upregulated genes against -log10 plot (Fig. 7B); and biological processes such as innate immune response, signal transduction, cell differentiation and proliferation were significantly enriched, indicating a broad impact on cellular functions (Fig. 7C).

**Figure 7.**
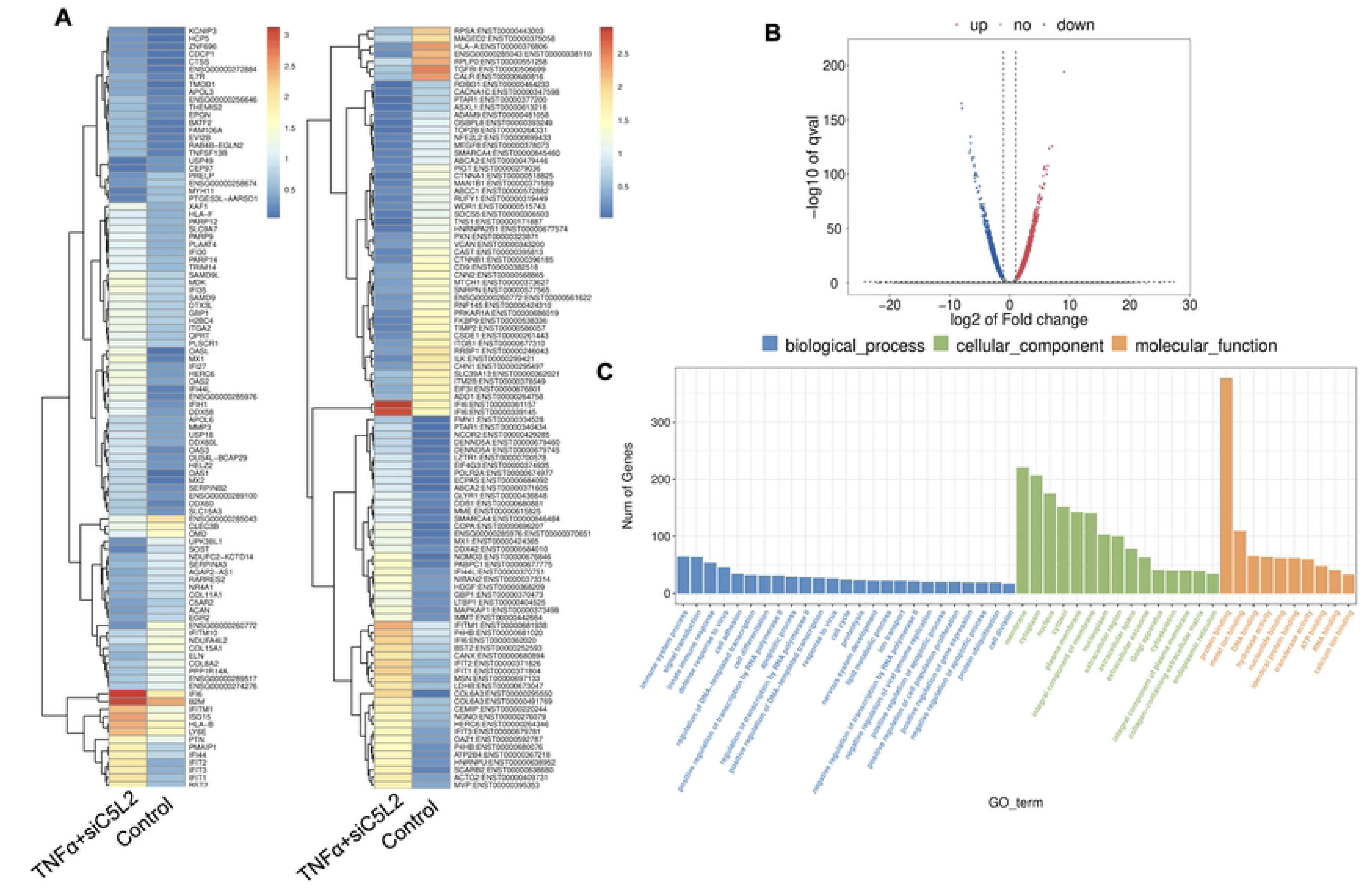
Analysis of differentially expressed genes (DEGs) in RNA-Seq. data. in TNFα and siC5L2 combined groups. **(A**) Heatmap of the differentially expressed genes between untreated or control cells and TNFα treated C5L2 silenced hDPSCs in dentinogenic media. Red and yellow stripes in the figure represent high expression genes, while blue stripes represent low expression genes. **(B)** Volcano map of differentially expressed genes (DEGs) between control cells and siC5L2 hDPSCs in dentinogenic media. The x-axis is the log2 scale of the fold change of gene expression in hDPSCs (log2(fold change)). Negative values indicate downregulation; positive values indicate upregulation. The y-axis is the minus log10 scale of the adjusted p values (–log10), which indicate the significant level of expression difference. The blue dots represent significantly upregulated genes with at least twofold change, while the red dots represent significantly downregulated genes with at least twofold change. **(C)** Significant enriched Gene Ontology (GO) terms among control and TNFα+siC5L2 treated hDPSCs based on their functions.

The primary GO terms identified among the genes include innate immune response and signal transduction under Biological Process (BP); integral component of membrane and cytoplasm under Cellular Component (CC); and protein folding and metal ion binding under Molecular Function (MF) [Figure 7. C]. COL6A and Interferon alpha-inducible protein 6 (IFI6). COL6A plays a crucial role in reparative dentinogenesis by contributing to the formation and organization of the extracellular matrix, supporting odontoblast differentiation and enhancing the tissue’s ability to respond to injury and mechanical stress (49). Its involvement in these processes makes it an important factor in maintaining the health and integrity of dental tissues, particularly following injury. IFI6 modulates the innate immune response by interacting with components of the retinoic acid-inducible gene I (RIG-I) signaling pathway, maintain a balance in the immune response, preventing overactivation that could lead to excessive inflammation or tissue damage (50).

Similarly, as mentioned above, twenty significant GO enrichment groups were identified including several enriched pathways related to regulation of immune response, collagen containing extracellular matrix and structural constituent, and, response to bacterium (Fig. 8A-B), that contribute to regeneration process (39). Sashimi plots for quantitative visualization of RNA sequencing read alignments revealed the number of variants and genomic mutation on chr3, chr11, Chr10 and chr19 in TNFα treated C5L2 silenced DPSCs in dentinogenic media against control (Fig. 8C).

**Figure 8.**
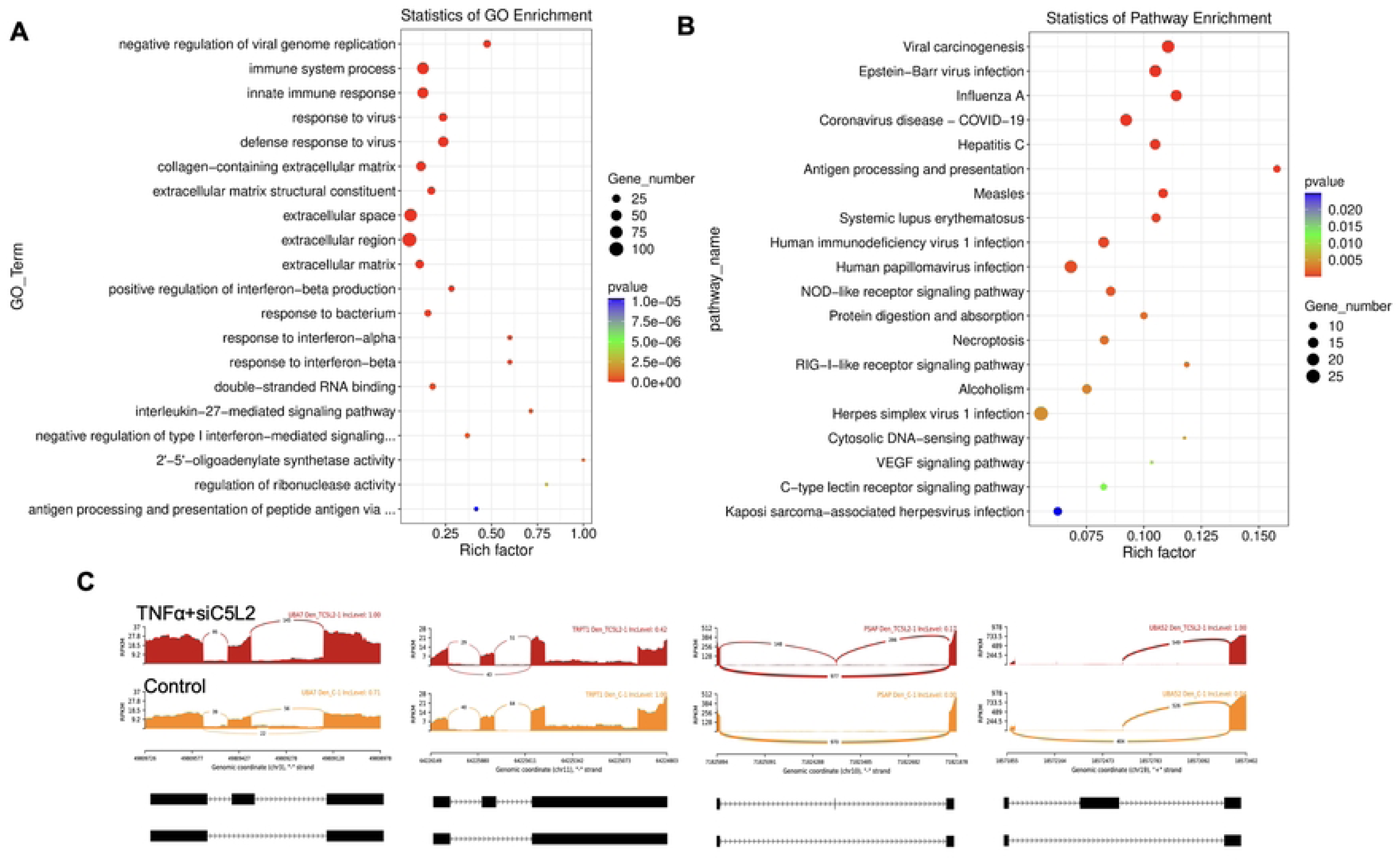
The top GO terms in the enrichment analysis among biological process, cellular component and molecular function (MF) terms in the enrichment analysis in RNA-Seq. data. in TNFα and siC5L2 combined groups. Significant enriched GO terms can be found in TNFα treated C5L2 silenced hDPSCs in dentinogenic media based on their functions compared with control. **(A-B)** Go-term and Reactome enrichment pathway analysis of up- and down-regulated DEGs. Dot plot shows top enriched Reactome pathways. The size of the dot is based on gene count enriched in the pathway, and the color of the dot shows the pathway enrichment significance. **(C)** Sashimi plots for quantitative visualization of RNA sequencing read alignments. Data were examined on sashimi plots where it revealed the number of variants and genomic mutation on chr3, chr11, Chr10 and chr19 in TNFα treated C5L2 silenced DPSCs in dentinogenic media against control. Red sashimi plots showing variants in TNFα+siC5L2 treated group and orange shows in control. While lower black annotations are Read alignments of alternative isoforms and genomic region of interest.

In the Chr3, showed strong splice junction connecting two exons with a high inclusion level of 1.00 in TNFα treated C5L2 silenced DPSCs, however in control also have a splice junction, but it connects different set of exons, with an inclusion level of 0.71. At the locus on chromosome 10, there is a large loop indicating an exon-skipping event, with an inclusion level of 0.17 in TNFα-treated C5L2-silenced DPSC in dentinogenic media, while the control groups show inclusion level at 0.00, indicating that this exon is completely skipped. For the locus on chromosome 19, control cells have very low inclusion level (0.04), suggesting that this exon is almost entirely skipped, whereas TNFα-treated C5L2-silenced DPSC a high inclusion level of 1.00, indicating that this exon is fully included in the transcripts. Overall, these results highlight how TNFα treatment combined with siC5L2 treatment significantly alters the splicing patterns of the genes shown, leading to different transcript isoforms compared to the control cells. Similarly, TCF/LEF transcription factors expressions increased (Fig. 9).

**Figure 9.**
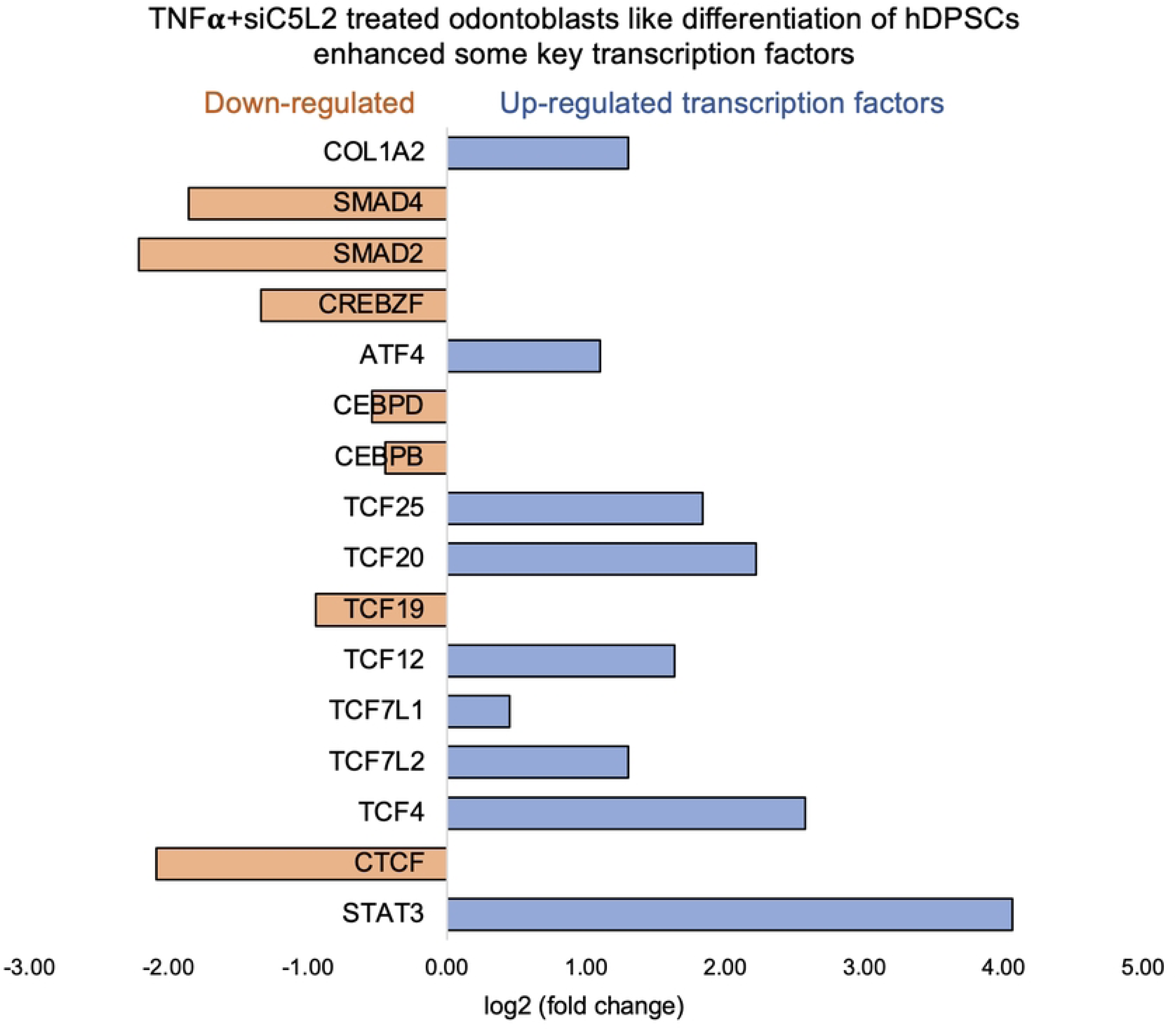
**Signaling pathways affected by C5L2 silencing in odontoblasts like differentiation of hDPSCs in dentinogenic media stimulated by TNFα**. hDPSCs were cultured and treated with or without TNFα (twice a week) in C5L2 silenced DPSCs and differentiated for 7 days. Cell lysates were collected, and RNA were prepared using RNeasy mini kit (Qiagen), and next-generation RNA sequencing was done using poly-A-RNA sequencing technique. Histogram showing upregulated and activated transcription factors (blue) and repressed or down-regulated transcription factors (orange). It is noteworthy that TCF (4, 12, 20, and 25) especially TCF4 highly up-regulated in C5L2 silenced TNFα-stimulated odontoblasts like differentiated DPSCs.

### 3.5. Real-time PCR confirms the expression of TCF family transcription factors and dentinogenic markers enhancement

We further evaluated the expression of various transcription factors and genes from sequencing by real-time PCR, and our result confirmed the expression of the TCF family transcription factors and other genes involved in dentinogenesis (Fig. 10).

**Figure 10.**
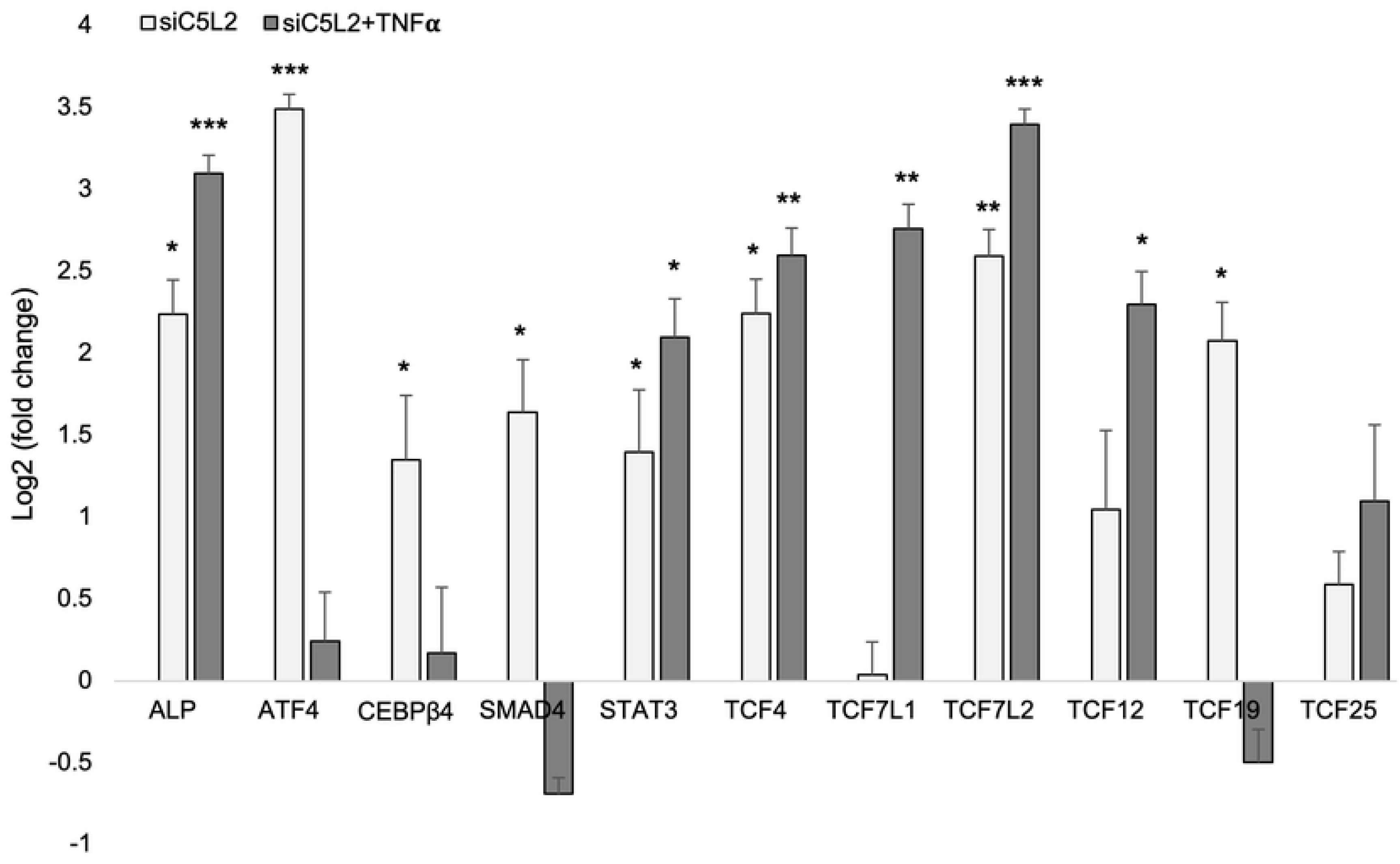
mRNA expression of TCF family transcription factors and other genes. mRNA expression during the odontogenic differentiation was quantified by real-time PCR. The elevated level of these markers represents odontoblast-like differentiation of DPSCs. * p < 0.05, ** p < 0.01 and *** p < 0.001 vs control fold change. The bar graph shows the mean ± SD of at least three independent experiments (n = 3) in duplicate.

## 4. Discussion

It is widely recognized that the regeneration of the dental-pulp complex is closely tied to inflammation. In caries, bacteria and their toxins affect odontoblast (51), while extension of bacterial infection to dentin affects the behavior of DPSCs in regeneration (52). Understanding the impact of inflammatory responses on DPSCs is crucial for uncovering the mechanisms in dentin and pulp regeneration. Due to the limited data available, this study takes a comprehensive approach to reveal this complex process and to identify key downstream molecules. Our data indicate that after inflammatory stimulation, DPSCs show significant changes in distinct gene expression profiles and transcriptomic alterations during odontoblastic differentiation. These include DPSS (3-fold increase), DMP-1 (4.1-fold increase), and alkaline phosphatase (ALP, 7.8-fold increase), indicating the critical role of inflammation in dentinogenesis.

Inflammation and immune responses are modulated by numerous pro-inflammatory cytokines including TNFα which acts through various pathways. The pro-inflammatory effects of this cytokine are well documented (53) and are involved in many osteo-immunological diseases including rheumatoid arthritis (54). Previous studies have proven the positive effect of low concentration of TNFα on increased osteogenic differentiation by upregulation od RUNX-2, ALP and OCN levels (55, 56). This dual effect of TNFα on osteogenic differentiation is directly related to its concentration, cell type and exposure time (10). In our current study, the low concentration and minimal exposure of TNFα on odontoblast like differentiating DPSCs for a specific time has increased the expression of osteogenic markers such as RUNX-2, ALP, DMP-1 and DSPP while also increasing the downstream transcriptional factors like LEF/TCF and CEBP-β. β-catenin and LEF/TCF form a complex with SMAD4 (42, 43) and, are needed to activate osteoblast gene expression and promote full differentiation. LEF/TCF and nuclear β-catenin can also act independently of each other by interacting with other transcription factors but the majority of β-catenin binds to chromatin via LEF/TCFs (57, 58). Among TCF family, particular attention is given to the role of TCF7L2, for having a central role in neural stem cell differentiation and postmitotic differentiation of some brain regions, impairments of which might contribute to mental disorders (44). The specific roles of LEF1, TCF7L2, and TCF7L1 in hippocampal developmental neurogenesis have been investigated in gene knockout studies, suggesting a redundant role for LEF1 and other LEF1/TCF family members. TCF7L1 knockout did not result in any hippocampal alterations, but the size of both the granule and pyramidal cell layers of the hippocampus was markedly reduced in TCF7L2 knockout mice (46, 59). In our results, C5L2 silencing showed significantly enhanced expression of TCF7L2, indicating role of C5L2 silencing in neural regeneration process and odontoblastic differentiation of DPSCs.

The STAT, SMAD, and ATF transcription factors play pivotal roles in inflammation-induced regeneration, particularly in processes like dentinogenesis and tissue repair. STAT proteins, activated through the JAK-STAT pathway, are critical mediators of cellular immunity, proliferation, apoptosis, and differentiation. In the context of dentinogenesis, STATs promote the transcription of genes essential for stem cell proliferation and differentiation, thereby contributing to the regeneration of dental tissues (48). SMAD proteins, which are key mediators of the TGF-β signaling pathway, are crucial for osteoblast and odontoblast differentiation. However, inflammation-induced factors like TNFα can inhibit SMAD signaling by activating NF-κB, leading to impaired differentiation and mineralization. This disruption can adversely affect dentinogenesis and bone formation, making the regulation of SMAD activity a potential therapeutic target for enhancing dental and bone regeneration (33). ATF genes, particularly ATF7ip, have been identified as negative regulators of osteoblast differentiation by influencing transcription factors such as Sp7. Additionally, ATF7ip plays a role in modulating immune responses, which can have significant implications for inflammation-driven regenerative processes. Targeting these transcription factors could, therefore, offer new strategies to modulate inflammation, enhance tissue regeneration, and improve outcomes in dental and bone repair therapies (34). These pathways collectively underscore the complex interplay between inflammation, transcriptional regulation, and tissue regeneration, highlighting the importance of fine-tuning these molecular signals to promote effective dentinogenesis and regeneration.

Absence of biomarkers in early detection and drug resistance are principal causes of treatment failure in various disease conditions. RNA-Seq has expanded the knowledge of molecular level pathologies progression and novel transcriptional changes in tissues. RNA-Seq technology has emerged as a powerful tool for identifying functional genes and pathways in diagnosing various disease mechanisms (60). It has discrete advantages over former methodologies and has transformed our understanding of the complex and dynamic nature of the transcriptome. It provides a more detailed, unbiased and quantitative view of gene expression, alternative splicing, and allele-specific expression. Recent advancements in the RNA-Seq workflow, from sample preparation to sequencing platforms to bioinformatic data analysis, has enabled deep profiling of the transcriptome and the opportunity to explore various physiological and pathological conditions (60, 61). However, available methods were having various limitations and restraints: the obligation for a prior knowledge of the sequences; problematic cross-hybridization artifacts in the analysis of highly similar sequences; and limited ability to accurately quantify lowly expressed and very highly expressed genes (61–63).

Stem cell therapy is a newly proposed technique for several diseases and human DPSCs are under consideration in regenerative medicine since last decade. DPSCs have a high capacity of differentiation and regeneration due its multipotency. Previous studies have described the differentiation potential of DPSCs including odontogenic, osteogenic, and neurogenic potential (64, 65), and DPSCs have been successfully used in bone tissue engineering (66). BM-MSCs are multipotent stem cells characterized by self-renewal and multilineage differentiation while DPSCs display MSCs-like characteristics but with easy accessibility, noninvasive isolation, limited ethical concerns, and high proliferation capacity. DPSCs are thought to be promising stem cell sources for clinical use (67). Yamada et al., (67) has summarized the regenerative capabilities of DPSCs in various systemic diseases, especially with regard to neural regenerative capacity compared to BM-MSCs, and concluded that DPSCs have comparatively much greater therapeutic potential. In our previous studies we have used human DPSCs to study odontoblast like differentiation and their effects on dentinogenesis (32, 36, 68) and found their suitability to study them in osteogenic differentiation studies.

Complement system is an early response to tissue damage and a critical component of innate immunity and inflammation (69, 70). This could be observed during bacterial infection and caries onset including the recruitment of immune cells due to the production of anaphylatoxin C5a and some opsonins (71). Beyond its role in immunity, the complement system participates in tissue regeneration (72). Another study has explained the linkage between inflammation and dental tissue regeneration through complement activation (73) and between inflammation and bone regeneration in BM-MSCs (74). It is noteworthy that complement receptor C5aR1 was also increased with treatment of TNFα indicating the complement C5a activation and involvement in the dentinogenic process and osteogenic differentiation of DPSCs confirming with our previous studies on DPSCs mediated stem cell therapy and regeneration process (19, 68). On the other hand, C5aR2 (C5L2) was not affected or reduced a little, while bacterial defense response was activated in siC5L2 treated group showing its positive effect in carious ailment. Our RNA profiling data indicate C5L2’s critical roles in governing the signaling pathway during inflammatory dentinogenesis. Previously, we have been working on C5L2 receptor and its involvement in inflammation mediated dentinogenesis (12, 36). Together with the current study’s outcome, the evidence of indirect involvement of C5L2 in regulation of dentin reparative process via BDNF/TrkB regulation and DSPP and DMP-1 increment, and up-regulation of downstream signaling through a key transcriptional factor i.e., TCF7L12 and β-catenin, proposed its potential as a target candidate to be considered to explore further in the field of stem cell based regenerative therapy. As C5L2 is still considered as a controversial receptor (also known to work against C5aR1), and there’s limited information available (75). We continue to explore its mechanistic aspects in the field of regenerative medicine.

## 5. Conclusion

Taken together, bulk RNA analysis revealed the key transcription factors and differentially expressed genes involved in inflammation induced odontoblast like differentiating DPSCs indicating critical role of TNFα and complement C5a receptor 2 silencing. This may direct future studies on specific transcription factors and role of TNFα induced inflammation in stem cell mediated regeneration therapies.

## Author contributions

Irfan M and Chung S contributed to conception, design, data acquisition, analysis, and interpretation, drafted and critically revised the manuscript. Irfan M conducted major experiments and contributed to data acquisition, analysis of the experiment; while Kim JH and Sreekumar SK conducted patrial experiments and contributed to data acquisition, analysis of the experiment and revised the manuscript. Chung S designed the original concept and contributed to data acquisition and interpretation and financially supported the project.

## Acknowledgments

The manuscript has been read and approved by all authors; and that all authors agree to the submission of the manuscript to *PlosOne*.

## Data availability

The datasets generated during and/or analyzed during the current study are available from the corresponding author on reasonable request.

